# Neuromodulation Influences Synchronization and Intrinsic Read-out

**DOI:** 10.1101/251801

**Authors:** Gabriele Scheler

**Affiliations:** Carl Correns Foundation for Mathematical Biology 1030 Judson Dr., Mountain View, Ca 94040

## Abstract

The roles of neuromodulation in a neural network, such as in a cortical microcolumn, are still incompletely understood. Neuromodulation influences neural processing by presynaptic and postsynaptic regulation of synaptic efficacy. Synaptic efficacy modulation can be an effective way to rapidly alter network density and topology. We show that altering network topology, together with density, will affect its synchronization. Fast synaptic efficacy modulation may therefore influence the amount of correlated spiking in a network. Neuromodulation also affects ion channel regulation for intrinsic excitability, which alters the neuron’s activation function. We show that synchronization in a network influences the read-out of these intrinsic properties. Highly synchronous input drives neurons, such that differences in intrinsic properties disappear, while asynchronous input lets intrinsic properties determine output behavior. Thus, altering network topology can alter the balance between intrinsically vs. synaptically driven network activity. We conclude that neuromodulation may allow a network to shift between a more synchronized transmission mode and a more asynchronous intrinsic read-out mode.

## Introduction

In this paper we present a realistic network model, akin to a cortical microcolumn [7, 8, 4, 25], and investigate its properties under the assumption of fast synaptic and intrinsic modulation as evidenced by neuromodulation [34]. We hypothesize that rapid synaptic efficacy changes allow to operate with different network topologies, and that network topology is a decisive factor towards creating and sustaining synchronized inputs vs. producing asynchronous input.

We have previously shown for a conductance-based neural model of striatal medium spiny neurons that neuronal heterogeneity expressed by the contribution of individual ion channels (such as delayed rectifier potassium channels or GIRK channels) may still result in uniform responses, if the neurons are driven with highly correlated synaptic input. If the same neurons are driven by more asynchronous, distributed synaptic input, the heterogeneity is manifest in the response patterns, i.e. the spike rates and the timing of the spikes (see [35]). These results were achieved using conductance-based point neurons [35]. Here we use two-dimensional neural models [17] to further investigate the effect and determine its significance in the context of a cortical neural network.

Due to Hebbian learning [36, 23], under normal conditions synaptic weights follow a lognormal distribution, which results in graphs with a heavy tail degree distribution (Fig. 8). Degree modification by rapid synaptic efficacy changes would allow to alter the density, but also the topology of the connecting graph. In this paper we examine the hypothesis that such changes in network topology actually occur, driven by neuromodulatory effects on presynaptic release or postsynaptic response ([41, 30, 22, 34]). We analyze this situation with two example graphs, and we also perform further analysis (Fig. 11) to show that there is a continuum of graphs which can be reached by rapid synaptic changes.

**FIG. 1.**
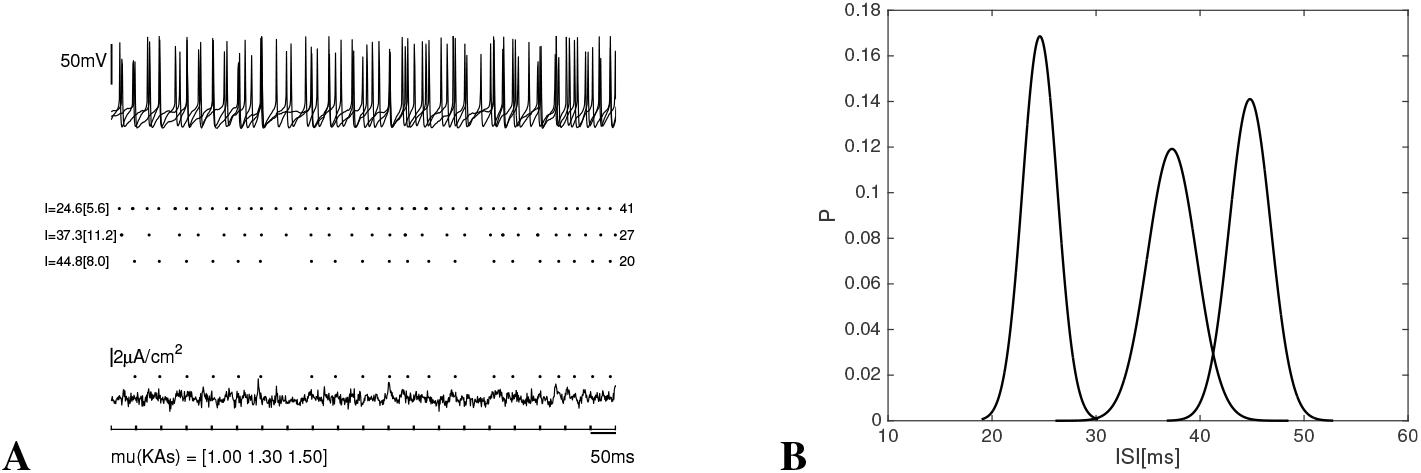
**A**. Frequency response of 3 conductance-based MSN model neurons with variable scaling of IAs to uncorrelated input **B**. Probability distributions of ISIs. We see a clear separation of frequency responses.

## Results

### Conditional Expression of Intrinsic Excitability in Conductance-based Models

We show how we can model gain as a stored intrinsic property, defined as a spike rate in response to constant or fluctuating input of fixed strength (A/Hz). We use a full ion channel based model (the MSN model [35]), with variation in the slow A-type potassium channel (IAs).

In Fig. 1A, we show the response of MSN model neurons with a scaling of *μ*_IAs_ = 10, 1.3, 1.5 to a noisy signal, derived from uncorrelated Poisson-distributed synaptic input. The top panel shows the development of the membrane potential, *V_m_*, over time for all neurons. The middle panel shows the spike-train for each neuron with the mean ISI and its standard deviation; the total number of spikes is shown on the right. The total number of spikes includes bursts, which were excluded from ISI calculation. The bottom panel shows the synaptic input. The dots correspond to the spiking events for a single neuron 3 (*μ*_IAs_ = 15). The resulting mean ISIs are 25, 37, and 45 ms. With a standard deviation of 6, 11, and 8, they are clearly distinguishable. This is also shown by the Gaussian distribution for the mean ISIs for each neuron type (Fig. 1B).

This model shows frequency-specificity as read-out of the relative contribution of the slow A-type potassium channel, indicated by the scaling factor *μ*_IAs_. The relative contribution of an ion channel corresponds to its density or distribution on the somato-dendritic membrane, or in some cases its specific localization at dendritic branch points. Experimental evidence has shown that this is a plastic feature for neurons.

We then employ highly correlated synaptic input, defined as in [35] (see *Materials and Methods*). We stimulate the same neurons with the correlated input and observe the spike pattern (Fig. 2). We can show that the frequency-specificity of the neuron disappears. Instead we see a time-locked spike pattern which is expressed by a similar spike frequency (Fig. 2A) and an overlap of the mean ISIs (Fig. 2B).

**FIG. 2.**
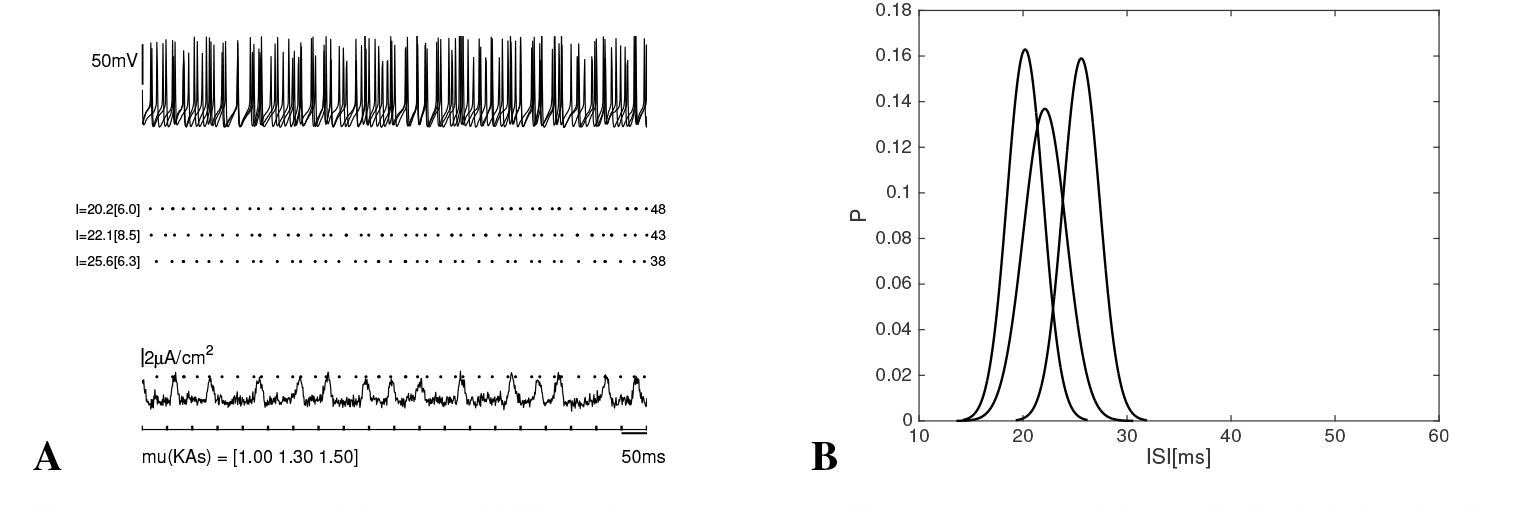
**A**. Frequency response of the same MSN model neurons as in Fig. 1 to correlated input. **B**. Probability distribution of ISIs. We see overlapping offrequency responses.

What this experiment showed is that a stored intrinsic property, the gain, is available to the processing network in a conditional manner. The property is continually expressed, the differences in ion channel density persist. Depending on the mode of stimulation, however, this property is manifested as intrinsic gain, or it is obscured when a neuron is driven by strongly correlated input.

### Results for Simplified Model Neurons

To continue with exploring this property of model neurons in a networked context, we switched to a simplified model neuron [17] and created a set of variations for this model (see *Materials and Methods*). We show the response of two-dimensional model neurons to asynchronous input in Fig. 3, and to regular, synchronous input in Fig. 4. In the first case, we have clearly separated frequencies, and in the second case, the ISIs are nearly identical with a narrow distribution. When we stimulate the neurons with irregular, but synchronous input, the ISIs become identical, but with a wider distribution to reflect the different duration of pauses between the synchronous stimulation (Fig. 5).

**FIG. 3.**
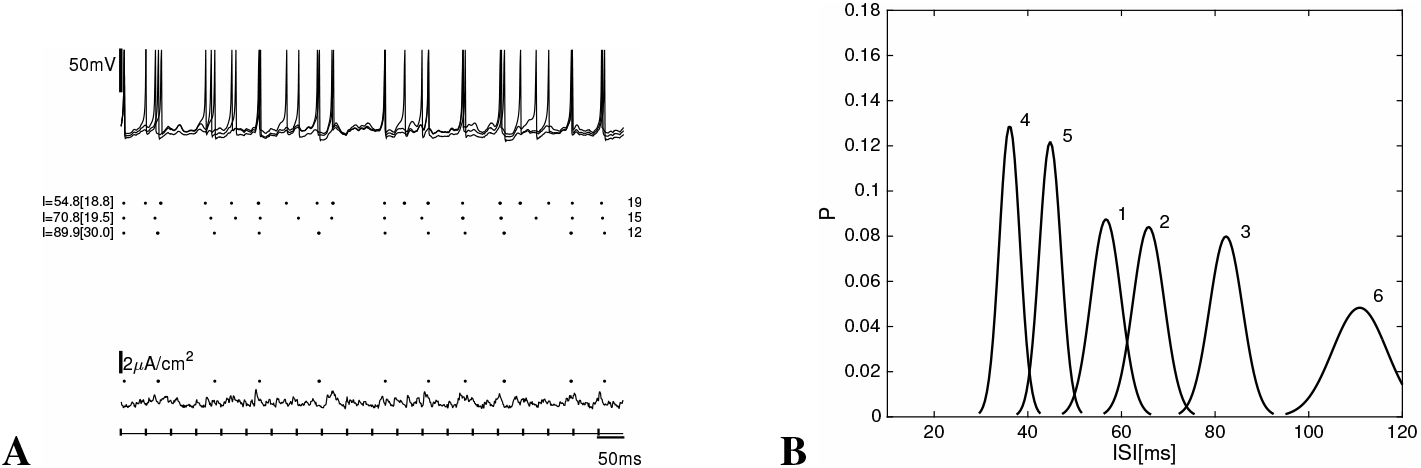
**A**. Spike response of the two-dimensional dynamic model neurons (1,2,3) to asynchronous input. **B**. Gaussian distributions of ISIs for six model neurons (1,2,3,4,5,6) to asynchronous input as in A. We see a clear separation of frequency responses for model neurons.

**FIG. 4.**
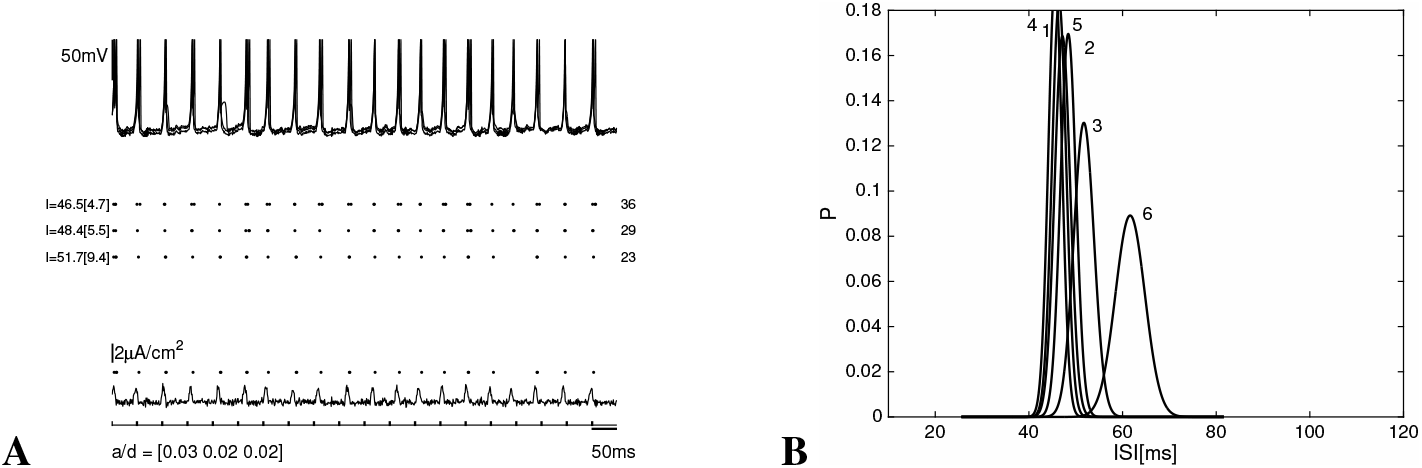
A. Spike response of the two-dimensional dynamic model neurons (1,2,3) to regularly timed, correlated input. **B**. Distributions of ISIs for six model neurons (1,2,3,4,5,6) to the same input as in A. We see strong overlapping of frequency responses, at about 50ms ISI, in accordance with the input. We notice that neuron 6 fires at lower frequencies than the input, it probably has a longer reset period, as seen in Fig. 3B.

**FIG. 5.**
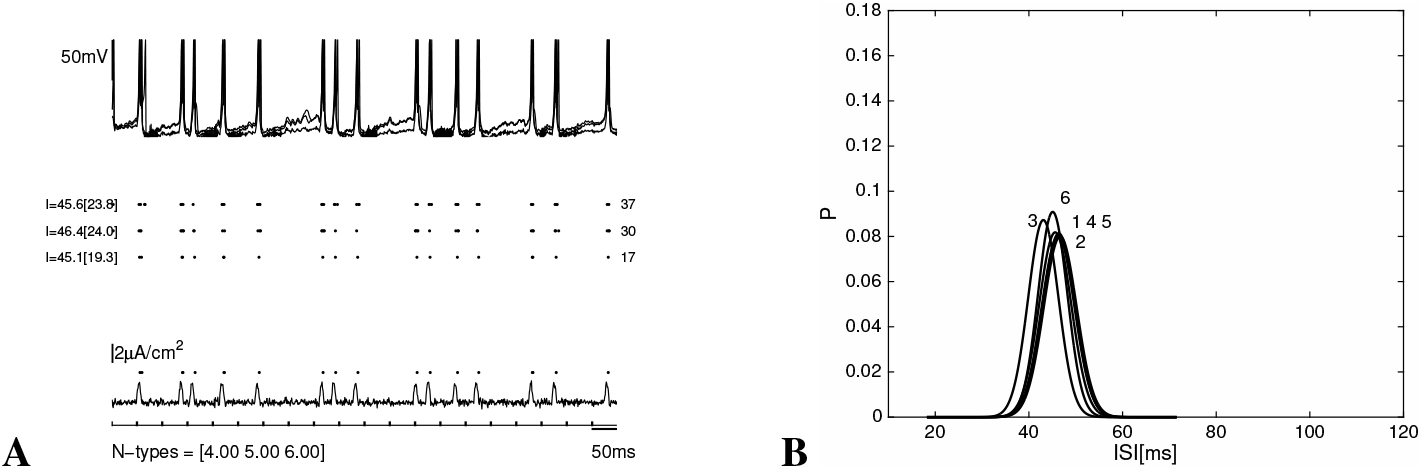
**A**. Spike response of the two-dimensional dynamic model neurons (1,2,3) to irregularly timed, correlated input. **B**. Gaussian distributions of ISIs for six model neurons (1,2,3,4,5,6). We see strong overlap offrequencyresponses.

### Multiplexing synchronous and asynchronous input

We may also consider the question of whether a neuron can simultaneously respond to an input and read out its stored spike frequency. If there are single synchronous events, which interrupt ongoing spiking, can we recover the intrinsic properties for each neuron? In Fig. 6 it is shown that this is possible. Fig. 6A shows the input and the synchronous responses, and in Fig. 6B we still see a clear separation of frequencies.

**FIG. 6.**
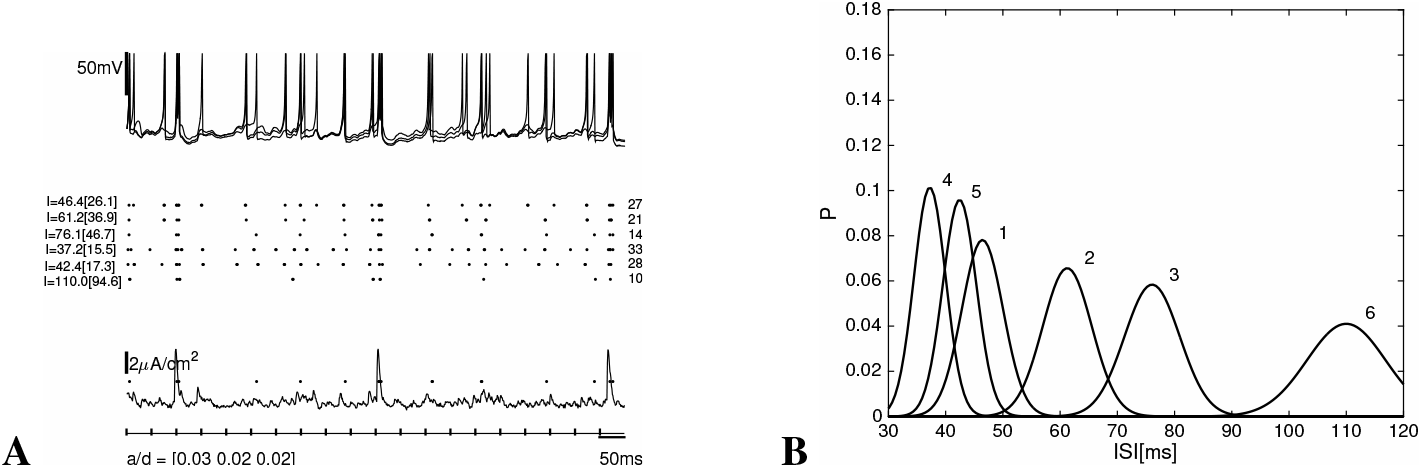
**A**. Multiplexed input and response of different model neurons (1-6). Three synchronous events are clearly represented in the spike pattern of all neurons. **B**. Mean ISIs for each neuron type. The separation of frequencies is kept. Compared to Fig. 3B the standard deviation is somewhat higher because of the additional spikes caused by strong synchronous input.

We conclude that we can multiplex asynchronous and synchronous input. It is also apparent that there needs to be a lower limit on the intervals between synchronous events that can be processed without disrupting intrinsic properties. This interval needs to be defined as functionally dependent on the intrinsic frequencies. In this case, it is 3/s for the synchronous events, with 10Hz for the slowest neuron.

### Synchronization depends on network topology

The simplified model neurons allow to create large networks of heterogeneous neurons and explore different topologies. We hypothesized that a lognormal graph, because of its hierarchical topology and the existence of hub neurons would lead to synchronization of action potentials - even with heterogeneous neurons - while a Gaussian topology would support asynchronous spiking behavior([2, 1]. We define synchronization *s* in a network by pairwise correlation (*Materials and Methods*). The spike frequency for each neuron type is assessed by the mean and standard deviation for ISIs, as before.

We first use a Gaussian connected graph (RG) with *N* = 210 and *K* = 1800 and use 7 different neuronal types (1-6, plus the generic neuron g) with 30 Neurons each (*Materials and Methods*). Fig. 7A shows an excerpt of the graph structure. We can see that the graph is connected such that all neurons have a comparable number of connections. This is also apparent in Fig. 8, where we can see a (narrow) normal distribution for connectivity for the Gaussian graph RG. Table 2 contains the graph characteristics for both graphs.

**FIG. 7.**
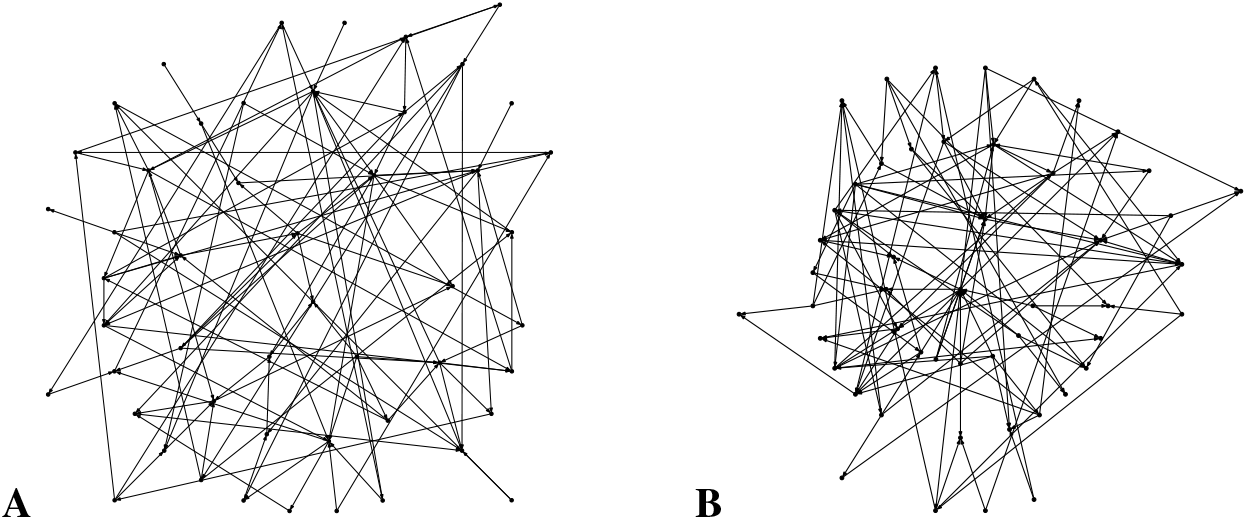
**A**. Part of a Gaussian Graph (RG), here for 50 neurons **B**. Part of a Lognormal Graph (LG1), for 50 neurons. The more regular, lattice-like structure of the Gaussian graph and the higher clustering and the appearance of highly connected ‘hub’ neurons in the lognormal graph is apparent.

**FIG. 8.**
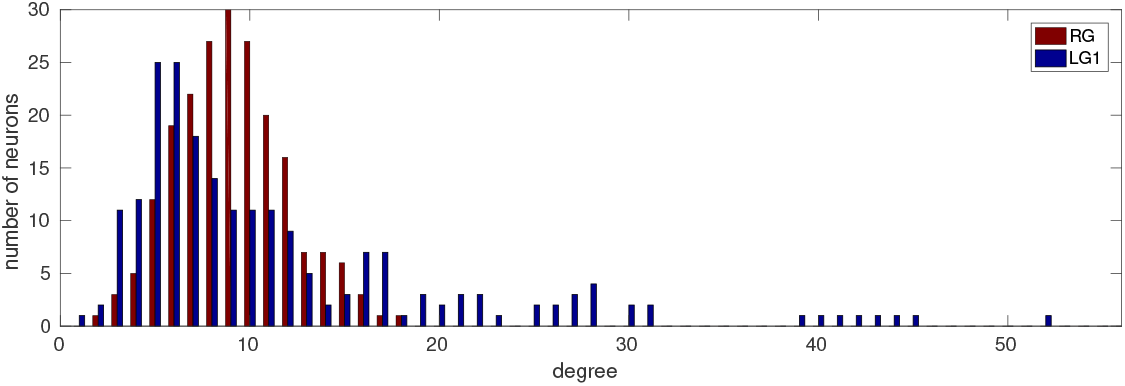
Degree histogram for the Gaussian Graph RG (red) and the lognormal Graph LG1 (blue). The LG has more neurons with
few connections. It also has a heavy tail of neurons with 20 and more connections (‘hubs’), which are lacking in the Gaussian graph.

*N* = 210 is about the size of a minicolumn or ensemble unit within a larger network with presumably dense interconnections [21]. The maximal density *d* = *K*/(*N* × *N* — 1) in a cortical microcircuit is estimated at 0.1 for 10^4^ neurons, 10^7^ synaptic connections, [26]. With ∼50% of synapses internal to the network, *d* = 0.04 — 0.07 is a realistic value for internal connectivity [21]. There is also a small background inhibition to all neurons present, implemented by 10% inhibitory neurons with Poisson-distributed firing and complete connectivity to excitatory neurons.

We now stimulate the graph by an initial stimulation to 10 excitatory neurons (for about 1s). In Fig. 9A, we see highly asynchronous neuronal activity after 1s of stimulation. The pairwise correlation values *s* is low (*s* = 0.11). Fig. 9B shows that each neuronal type retains its own frequency, i.e., has its own typical ISI, separated from other neuronal types. We also notice that some neurons fire with low frequencies (5Hz) and others with higher frequencies (20Hz). Very low firing neurons (2Hz) which are typical for cortex are not represented in this model.

**FIG. 9.**
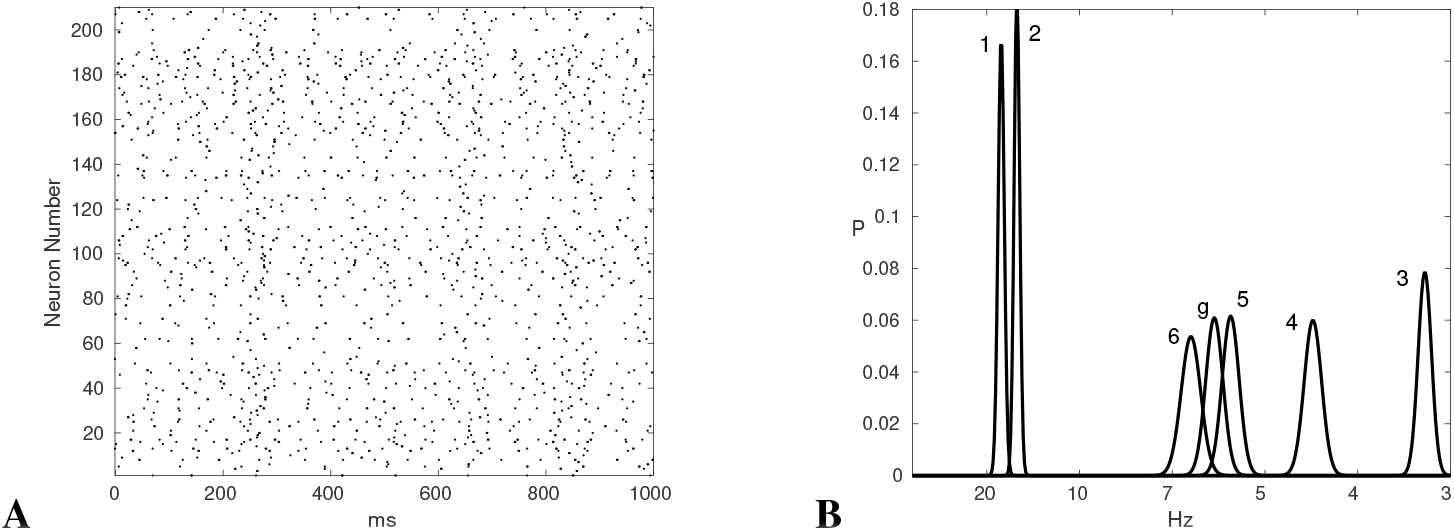
**A**. Asynchronous behavior in the Gaussian graph with variable neuron types. Groups of neuronal types are apparent in the rasterplot. Some structure is probably due to background inhibition. **B**. Average spikes/s with probability distributions for all neuronal types, with clear separation by frequency. Pairwise correlation is *s* = 0.11.

Next we changed the topology of the network to a lognormal graph (LG1), with *N* = 210 and *K* = 1924 and used the same neurons as before.

Fig. 7B shows an excerpt of a lognormal graph structure. The connectivity structure seems much denser, because of ‘hub’ neurons in the center of the graph. In Fig. 8, we can see a much wider distribution of degrees for the lognormal graph (blue), with a number of nodes with high connectivity. Presumably, those nodes are capable of synchronizing the network, because they can reach many neurons simultaneously. What is the effect in the presence of neural heterogeneity?

Fig. 10 shows that a high amount of synchronization can be achieved in spite of heterogeneity of intrinsic frequency of model neurons. The rasterplot (Fig. 10A) shows the activity in LG1 with the same neurons and the same stimulation as before. The overall correlation, defined by pairwise correlation of neurons, is much higher (s=0.32). The distribution of ISIs in this case is strongly overlapping (Fig. 10B), similar to Fig. 4B, where neurons were explicitly driven by highly synchronous input. This means that synchronization is dependent on the network topology, and a lognormal graph exhibits a higher tendency for pairwise synchronization. Also, that neuronal heterogeneity is apparent in an asynchronous network mode but is repressed in a synchronous firing mode.

**FIG. 10.**
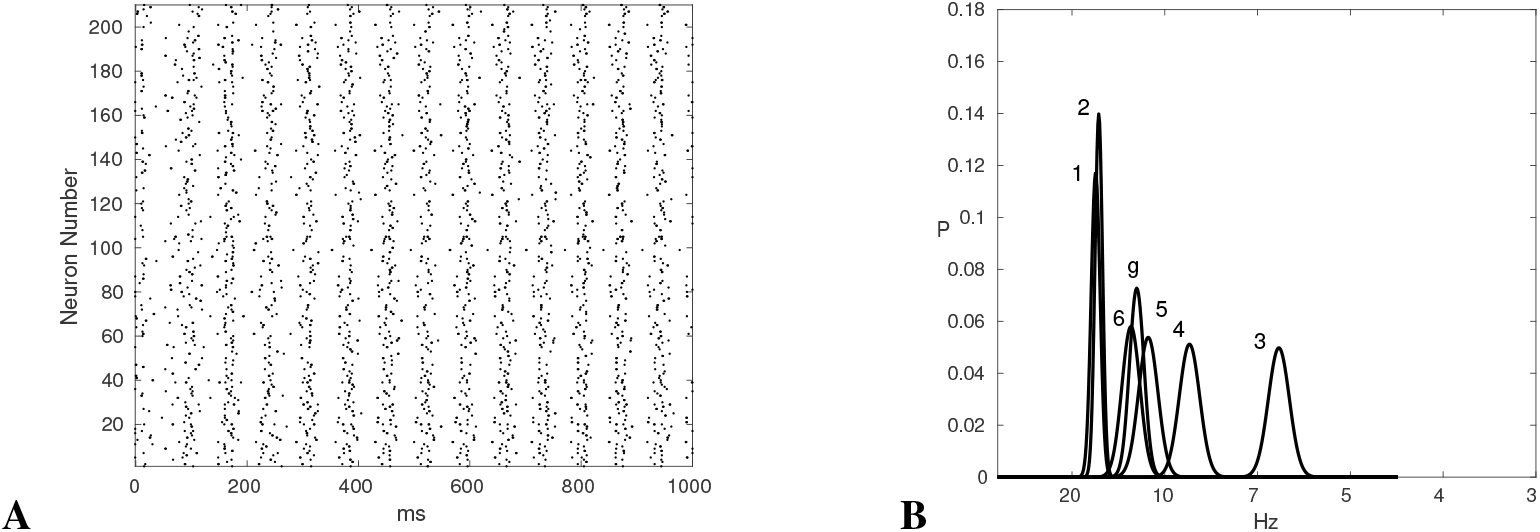
**A**. Synchronization in a heavy-tail graph (LG1) with variable neuron types. The rasterplot shows that different neuronal types respond uniformly. **B**. Frequency distributions. High overlap between neuronal types is apparent. Pairwise correlation in the graph is high with *s* = 0.32.

**FIG. 11.**
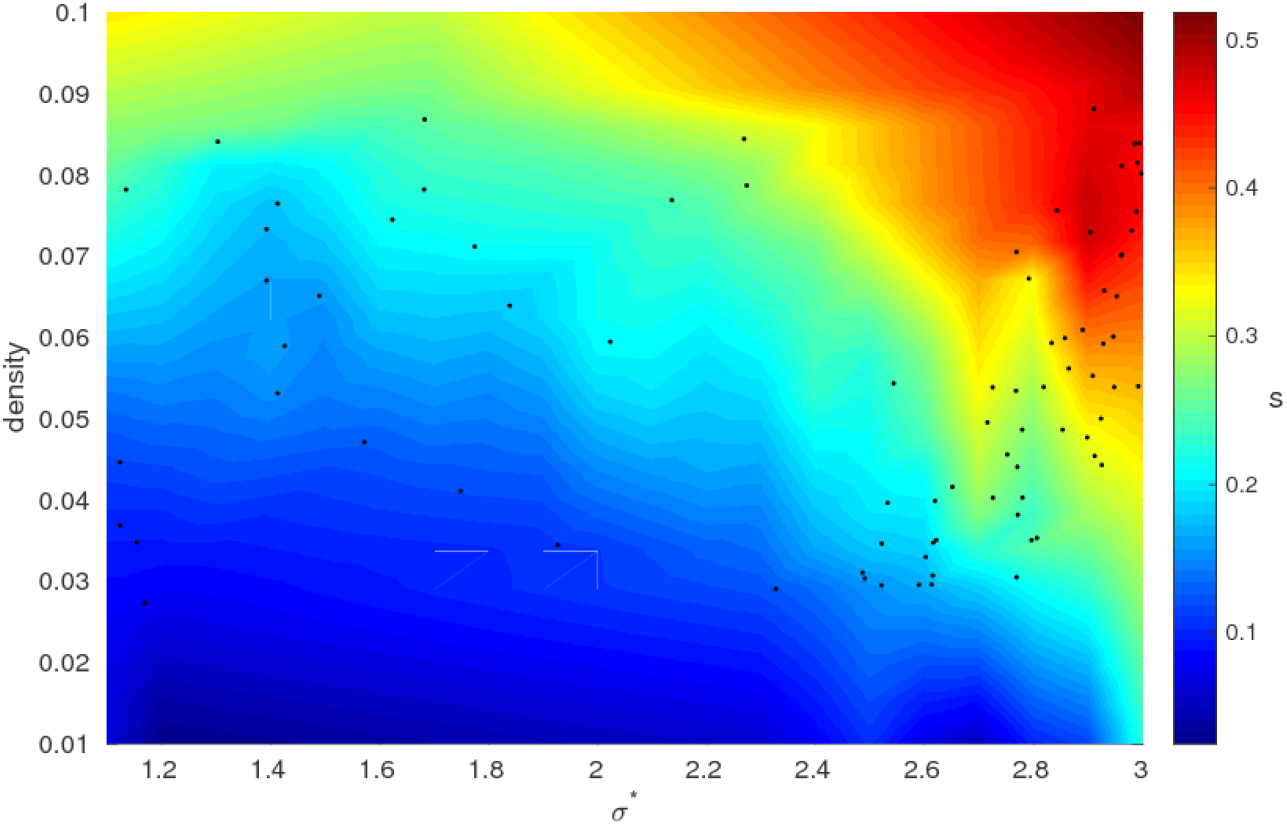
Synchronization *s* dependent of network topology: density and dispersion. The experimentally attested dispersion for weights in cortical tissue [36] is *σ*\* = 2 – 3.5, with a mean at 3. We achieve higher synchronization in the lognormal region, also dependent on density, but no synchronization in the Gaussian (low dispersion region) (*σ*\* < 1.5), except close to maximal connectivity. Black dots signify actual measurements. There seem to be no abrupt transitions.

### Dependence ofsynchronization on graph properties

We could show that differences in intrinsic properties appear or become more prominent when there is less synchronicity in a network. In our model, the pairwise synchronicity *s* is dominated by the network topology, more precisely by the width of the degree distribution (dispersion) ranging from Gaussian to lognormal with a heavy tail.

To confirm this observation we used a number of intermediate graphs and mapped the pairwise synchronization dependent on the dispersion *σ*\* (Fig. 11). The graphs RG and LG1 that we used have values of *σ*\* = 1.44 and *σ*\* = 2.89 (*Materials and Methods*). They have the same density, i.e., the same number of connections and neurons 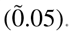. Additionally, we analysed the dependence of synchronicity on the density of the graph between 0.01 and 0.1 (Fig. 11).

There is higher synchronization in the lognormal region, especially with higher *σ*\* > 2.5 but no synchronization for Gaussian graphs. For heavy-tail graphs, synchronization depends linearly on the density between *d* = 0.03 - 0.08 (*s* = 0.2 - 0.5).

How are the different graphs related? We hypothesized that fast synaptic switching [37] by neuromodulation could change the network topology sufficiently to switch from a synchronous to an asynchronous regime. In Fig. 12, we plot the number of edges that were changed to achieve different dispersions *σ*\* of a graph. The algorithm used was a simple greedy algorithm (*Materials and Methods*), which is suboptimal, i.e. overestimates the number of edges required. It appears that 30-50% of edges changed would be sufficient.

**FIG. 12.**
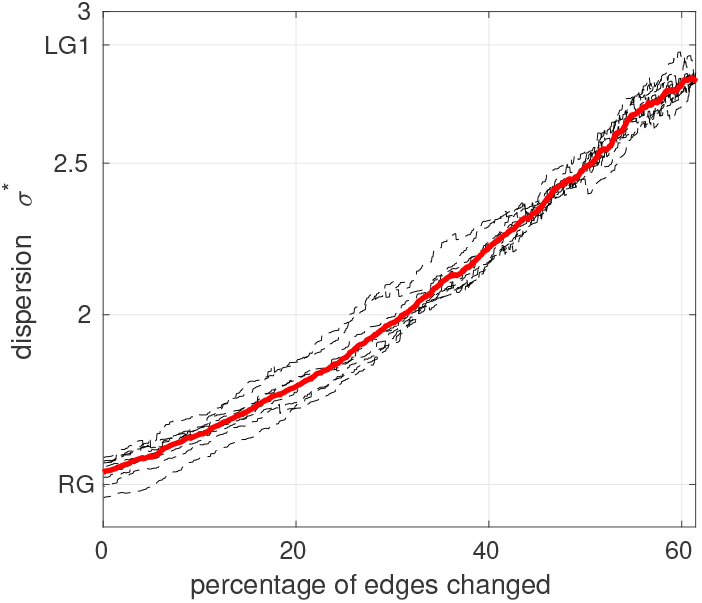
Transition between a lognormal graph and a Gaussian graph: For K=1900, N=210, d=0.43, mean over 10 trials, the percentage of edges changed to achieve a narrow degree distribution. The algorithm is not optimized (Materials and Methods) and overestimates the number of edges that have to be changed.

## Discussion

### Network Topology, Synchronization and Intrinsic Read-out

We employ a parameterizable two-dimensional neural oscillator model to encode different intrinsic excitability manifested by different frequency responses to constant input. What the experiments show is that a stored intrinsic property, the gain, is available to the processing network in a conditional manner: The gain is continually present, the differences in ion channel density persist. Depending on the mode of stimulation, however, this property is manifested as spike rate, or it is obscured when a neuron is driven by strongly correlated input. This is interesting because it shows a property of memory that synaptic plasticity lacks: the memory is not always ‘read-out’ in any processing step. It is conditional, it can be accessed or ignored depending on the state of the network. This seems to be an essential property of memory in any intelligent system.

Different statistical properties of synaptic input can be modeled by a variability in the correlation properties of input neurons. In a network model, this means that the overall correlation in the network determines what input a neuron receives. With a Gaussian degree distribution topology, correlation is low and neurons fire irregularly with their own preferred frequency. With a heavy-tailed, lognormal topology, correlation is higher, and neurons fire when they receive correlated input, irrespective of intrinsic properties. I.e. driving neurons by correlated vs. uncorrelated input leads to uniform spiking behavior vs. read-out of stored differences in ion channel conductances.

### Inhibition

A restriction of the present model with respect to a biological simulation model is the simplified treatment of inhibition. However, experimental work shows that cortical parvalbumin-expressing (PV+), fast-spiking interneurons have no connection specificity to pyramidal neurons, rather they present as an ‘unspecific, densely homogeneous matrix covering all nearby pyramidal cells’ ([31], p. 13260), which corresponds to our model.

Conditions for neuronal read-out may include the activity of inhibitory neurons. Inhibition and excitation are tightly linked by feedback interaction. [12] suggested that the close coupling of inhibition and excitation in cortical tissue cancels out purely input-dependent, i.e. not network generated synchrony. [14] suggested that with highly correlated input, both inhibitory and excitatory, the neuron may receive less input which allows it to be driven only by strong synaptic input, with distributed input it may receive a barrage of excitatory and inhibitory inputs where the membrane voltage remains close to firing threshold and the neuron fires continuously. In our sense, it is reading out stored properties. Both inhibitory and excitatory synaptic input will conform to be asynchronous or synchronous, to drive neurons or to causes a neuron to emit spikes according to its own stored intrinsic properties.

However, neuromodulation has effects on inhibitory neurons as well [24, 40], which we have not modeled. Further simulations will show, whether the I-E coupling is altered during enhanced neuromodulation, or whether the effects are synergistic to the present results.

### The Role of Neuromodulation

Neuromodulation influences both intrinsic properties and synaptic connectivity [34], e.g. acetylcholine, (via nucleus basalis stimulation), noradrenaline (via LC stimulation) or dopamine (via VTA stimulation) [13, 34]. Experimental estimates on the distribution of synaptic neuromodulatory receptors are at approximately 30%-50% of connections [37]. That is sufficient to transform the topological properties of a graph, such as the dispersion of its degree distribution from heavy-tailed graph to a more Gaussian, less clustered graph without requiring tight optimization for the positions of neuromodulatory receptors (Fig. 12).

Neuromodulation disables or enhances various ion channels, such as Sk-channels which guide reset times after a spike, or A-type potassium channels which influence latency to spike [35]. In this way, neuromodulation influences intrinsic properties. If neuromodulation reduces synchrony by acting at synaptic receptors, it uncovers intrinsic heterogeneity, and induces a mode of processing that allows read-out and storing of intrinsic properties. Depending on the neuromodulator used, and the amplitude and duration of the signal, different soma-dendritic ion channel profiles would emerge.

In the synchronous mode, intrinsic heterogeneities are reduced in the presence of tightly correlated input which drives neurons reliably. This invariance of neuronal intrinsic properties in synchronous mode allows synaptic transmission and information processing independent of neuronal heterogeneity.

The idea of introducing synchronous events by common input to an asynchronous background, and in this way use reliable synaptic transmission without affecting the state of the system (multiplexing) has also been documented in experimental results. For instance, ([15], Fig. 4A) shows a case of multiplexing in response to behavioral stimuli. In this case, intrinsic read-out can continue, and single events are transmitted reliably through driven activations.

Why should synchronization properties be switched by neuromodulation? Increased correlation in the network supports population-coded information to be propagated effectively [42]. Turning on neuromodulation would decorrelate an area and increase the capacity for information coding in an ensemble or a cortical microcolumn [39]. This area would become an information source to surrounding areas. When turned off, increased correlation would allow this area to transmit information and to disregard the stored neural memory.

### Relation to Experimental Evidence

Basal forebrain stimulation, which results in increased acetylcholine release and muscarinic/nicotinic receptor activation, decreases correlation between cortical neurons ([11, 9], [27] (Fig. 3.C)). Likewise, [28] (Fig. 3,4) shows reduced noise (internal) correlations with cholinergic stimulation, while inactivation of the basal forebrain caused more synchronized activity. [19] shows reduction of correlation for task-relevant perception, where presumably task-relevance causes neuromodulatory activity. [10] provides evidence for the involvement of noradrenaline in desynchronization of cortical state and the enhancement of sensory coding.

There is considerable evidence [16, 38, 33, 3] showing that several neuromodulators, including at least noradrenaline and acetylcholine, modulate pairwise spike correlation, such that strongly synchronized states (anesthesia, slow wave sleep) have high correlation and low neuromodulation, while asynchronous states (normal waking), with higher neuromodulation, have lower pairwise correlation.

[3] observed intrinsic fluctuations in synchronization of cortical networks during wakefulness which correlated with the amount of encoded perceptual information and perceptual performance. Their results showed a mean decrease in correlations from synchronized to desynchronized state corresponding to perceptual performance by approximately 20%, similar to values observed during attention [29], and after adaptation [14]. We have shown (Fig. 11) that correlation changes are continuous with network topology and a 20% correlation change is well within the range of the current simulations. Importantly, the results in [3] point to fluctuations in synchronization that reflect local changes in network activity rather than just global cortical state dynamics which have traditionally been associated with central neuromodulatory release.

The role of presynaptic neuromodulation in suppressing cortical connections [30, 22] and changing attractor states [20], as well as allowing rapid synaptic weight changes [37] has previously been assessed. Theoretical work has also emphasized the connection between correlations and information content [5, 39, 6, 32].

Here we bring these observations together to suggest that neuromodulation of synapses may alter network topology and in this way bring about an increased decorrelation of spiking, and a more asynchronous state, with a higher informational capacity. It may provide a general explanation (a) on how fluctuations in synchrony can be engineered rapidly and in small cortical areas and (b) why intrinsic memory may be conditional, accessible only at certain times and in a localized fashion.

## Conclusions

We created a number of different parameterized neuron models to capture neuronal heterogeneity. This affects the properties of the neuron such that it has less or more intrinsic excitability, leading to different firing rates when stimulated in an asynchronous way. Under synchronous stimulation the differences are greatly reduced.

We also suggested that synaptic neuromodulation can be an effective way of rapidly altering network topology. We investigated changes in network topology along the dimensions of Gaussian vs. heavy-tailed degree distributions. We hypothesized that heavy-tailed graphs produce more globally synchronized behavior than comparable Gaussian graphs. In accordance with the hypothesis, we find that in a heavy-tailed graph, because of high population synchrony, the difference between neuronal intrinsic properties is minimized, while a Gaussian graph allows read-out of neuronal intrinsic properties. Thus, altering network topology can alter the balance between intrinsically determined vs. synaptically driven network activity.

## Materials and Methods

### Conductance-based neuron model and synaptic input

The conductance-based neural model of a striatal medium spiny neuron is described in detail in [35]. The membrane voltage *V_m_* is modeled using the equation

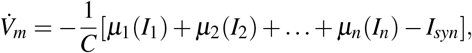

where the *I_i_* are the currents, induced by the individual ion channels. Variability of the neuron is modeled by modifications to *μ_i_*. This model includes ion channels for Na (INa), K (IK), slow A-type K channels (IAs), fast A-type K channels (IAf), inward rectifying K channels (IKir), L-type calcium channels (ICaL), and the leak current (I_leak_). The definition of all parameters and the dynamics of the ion channels can be found in [35].

For the experiments in this paper, we only focus on variability induced by changes in the strength of the slow A-type K channels. The total current contribution for this channel is *μ_IAs_* where *μ* was selected between 1.0 and 1.5.

In order to illustrate the variability in neuron behavior, we excited the neuron model by input signals, resembling two kinds of synaptic input: uncorrelated and correlated. These signals were generated by superposition of excitatory and inhibitory spikes from individual Poisson-distributed spike trains (50 excitatory and 10 inhibitory), and biased Gaussian background noise.

The amount of pairwise correlation in these spike trains governs the type of input signal. A high correlation factor was used in order to generate sequences which have short periods (10-15ms) of high activity.

### Heterogeneity in a two-dimensional model

In order to do large-scale simulation we needed to employ a simple, computationally tractable neuron model. We used a two-dimensional model of a neural oscillator (cf. [17]), and employed an instantiation of the model with parameters fitted to the general properties of cortical pyramidal neurons [18] as a generic model g. The model consists of an equation for the membrane model *v* (Eq. 1), fitted to experimental values for cortical pyramidal neurons, and an equation for a gating parameter *u* (Eq. 2).

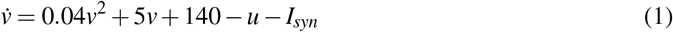

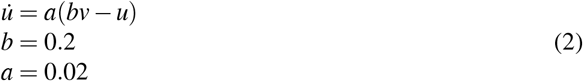

When the neuron fires a spike (defined as *v(t)* = 30*mV*), *v* is set back to a low membrane potential *v*: = *c*;*c* = -65.8*mV* and the gating variable u is increased by a fixed amount *d (u*:= *u* + *d*;*d* = 8) (cf. [18]). This formulation allows for a very simple neuron model, which avoids the explicit modeling of the downslope of the action potential, and rather resets the voltage. Time-dependence is modeled by the gating variable *u*.

Neuronal heterogeneity is achieved by systematic variation of inactivation parameters. By varying *d*, we can vary the inactivation dynamics of the model after a spike, by varying *a* we vary the inactivation dynamics throughout the computation. In this way, we can attempt to model neuronal variability in activation/inactivation dynamics, which is sufficient to model frequency-selectivity as a stored intrinsic property. The parameters used in this paper for different neuron types are listed in Table 1.

**Table 1.** Parameters for different neuron types

| name | $a$ | $b$ | $c$ | $d$ |
| --- | --- | --- | --- | --- |
| generic | 0.02 | 0.2 | -65 | 8 |
| type1 | 0.025 | 0.2 | -65 | 6 |
| type2 | 0.02 | 0.2 | -65 | 9 |
| type3 | 0.015 | 0.2 | -65 | 12 |
| type4 | 0.015 | 0.15 | -65 | 14 |
| type5 | 0.022 | 0.3 | -65 | 14 |
| type6 | 0.022 | 0.3 | -65 | 9.5 |

### Graph properties

We created graphs of *N* (= 210) excitatory neurons, and *K* 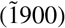 excitatory connections. For the excitatory neurons, we use randomly connected graphs (N,K) with different dispersion *σ*\*. This corresponds to normal (Gaussian) to lognormal graphs with different widths and length of the heavy tail. We model inhibition by Poisson-distributed inhibitory synaptic input directly onto excitatory neurons.

We use specific instantiations of these graphs (RG, LG1) for the simulations. Table 2 shows global graph characteristics for the Gaussian graph (RG), the lognormal graphs(LG1), and intermediate graphs LG2, LG3, LG4.

**Table 2.** Graph Properties for Excitatory Connections

| Property | RG | LG1 | LG2 | LG3 | LG4 |
| --- | --- | --- | --- | --- | --- |
| N | 210 | 210 | 210 | 210 | 210 |
| K | 1880 | 1924 | 1895 | 1920 | 2050 |
| d | 0.042 | 0.043 | 0.043 | 0.043 | 0.047 |
| indegree | 8.95[2...17] | 9.5[0...26] | 9[0...24] | 9.14[0...30] | 9.75 [0...25] |
| outdegree | 8.95[4...18] | 9.5[0...56] | 9[0...58] | 9.14[0...53] | 9.75 [0...54] |
| cluster index | 0.04 | 0.068 | 0.064 | 0.062 | 0.08 |
| mean plength | 2.66 | 2.41 | 3.7 | 3.72 | 2.32 |
| $\sigma^*$ | 1.44 | 2.89 | 2.5 | 2.71 | 2.71 |
| synchron s | 0.11 | 0.32 | 0.26 | 0.31 | 0.34 |

### Definitions

We define synchronization *s* in a network by pairwise correlations: for each neuron *n_i_*, we count, for each other neuron *n_j_* the number of spikes which occur within a window *W (W* = 10*ms*) of *n_i_*’s spike events, divided by the total number of spikes for *n_i_*. More precisely, for each neuron, we bin all firing events into 5ms bins. We then count the number of other neurons, which fire in a 10ms window around the (start) of the bin. The synchronization *s* is then the average over all neuron pairs in the network:

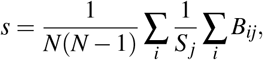

where *B_ij_* is the number of spikes that neurons *i* and *j* have in common within a moving window of *W* = 10*ms* during the entire measuring time. *S_j_* is the number of spikes of neuron *j* during the entire measuring time.

The rewiring algorithm used to change the properties of a graph G is a greedy algorithm, which iteratively selects the node *s* with the highest degree. One of its edges is then rewired to random nodes with lower degrees, decreasing *σ*\*. The algorithm terminates, when the value of *σ*\* falls below a given threshold.

## Availability of Data and Materials

Available at https://github.com/Scheler/CNeuroSyn.

